# Targeting catecholate siderophore systems enables precision depletion of gut enterobacteria during colitis

**DOI:** 10.64898/2026.10.02.756324

**Authors:** Grant J. Norton, Anna M. Dudek, Romana R. Gerner, Chuchu Guo, Vanessa Castillo, Felix A. Argueta, Connor J. Beebout, Ghazal Kamyabi, Julia Drushell, Gregory T. Walker, Chia-Yun Hsu, Shingo Bessho, Sean-Paul Nuccio, Hiutung Chu, Karsten Zengler, Elizabeth M. Nolan, Manuela Raffatellu

## Abstract

Adherent-invasive *Escherichia coli* (AIEC) is frequently isolated from the dysbiotic gut microbiome in inflammatory bowel disease and is implicated in disease progression, highlighting its potential as a therapeutic target. Here we show that iron acquisition is essential for AIEC colonization of the inflamed gut. Mechanistically, four catecholate siderophore receptors act redundantly to support AIEC growth under iron-limited conditions and during colitis. Immunization of colitis-prone *Il10*^-/-^ mice against catecholate siderophores reduced intestinal AIEC colonization, particularly its association with the mucosa, while largely preserving overall gut microbiome composition. Longitudinal metagenomic profiling identified immunization-associated shifts in *Akkermansiaceae* and *Lactobacillaceae*, and experiments in gnotobiotic *Il10*^-/-^ mice demonstrated that *Limosilactobacillus reuteri*, a member of the *Lactobacillaceae*, limited AIEC colonization, whereas *Akkermansiaceae* exacerbated disease. Our findings establish catecholate siderophore-mediated iron acquisition as a therapeutic target in AIEC and provide proof-of-principle that virulence-targeted immunization can selectively antagonize a disease-associated bacterium while preserving the gut microbiota.

## INTRODUCTION

Inflammatory bowel diseases (IBD) are chronic inflammatory disorders of the gastrointestinal tract with rapidly increasing global prevalence ^1–3^. Although the precise cause(s) of IBD remain unclear, host genetics, environmental factors, and an imbalanced gut microbiota (*i.e.*, dysbiosis) all contribute to disease onset and progression ^4–7^.

In the healthy gut, the microbiome is largely composed of obligate anaerobes of the phyla *Bacillota* (formerly *Firmicutes*) and *Bacteroidota,* which, in symbiosis with host epithelial cell metabolism, maintain the gut as an anoxic environment ^8,9^. In IBD patients, chronic inflammation of the gut mucosa alters the intestinal milieu and is associated with dramatic changes in the microbiota. Obligate anaerobes are most susceptible to host inflammatory responses, including the production of reactive oxygen and nitrogen species. The resulting disruption of intestinal hypoxia provides a fitness advantage to facultative anaerobes, such as *Enterobacteriaceae,* which thrive in the inflamed gut ^10–13^. In IBD, particularly Crohn’s disease (CD), the dysbiotic gut microbiome is often characterized by the presence of a specific *Escherichia coli* pathovar, adherent-invasive *E. coli* (AIEC), which is thought to exacerbate disease severity ^14–20^. Understanding the mechanisms that enable AIEC and other potentially harmful microbes to colonize the inflamed gut and interact with members of the gut microbiota is essential for developing new therapeutic strategies for IBD.

Iron is an essential metal nutrient that promotes the bloom of *Enterobacteriaceae* in the inflamed gut ^21^. In the host, iron is predominantly sequestered in intracellular compartments or bound to proteins to limit its reactivity and toxicity ^22^. As a result, the bioavailability of free iron is limited during homeostatic conditions, and is further reduced during inflammation and infection through a host defense mechanism known as nutritional immunity ^23^. To overcome this limitation, *Enterobacteriaceae* secrete siderophores, small molecules that scavenge ferric iron from the local environment and deliver it to the bacterial cell via substrate-specific cognate receptors on the bacterial outer membrane ^24–27^. Enterobactin (Ent) is a catecholate siderophore widely produced and secreted by commensal and pathogenic *Enterobacteriaceae* to acquire iron in the gut. To counteract this bacterial iron acquisition strategy, the host produces lipocalin-2, an antimicrobial protein that binds to and sequesters iron-laden Ent (reviewed in ^28^). The production of lipocalin-2 represents a highly effective antimicrobial defense against enteric microbes that depend exclusively on Ent for iron acquisition ^29^. However, many pathogenic *Enterobacteriaceae* (including *Salmonella*, uropathogenic *E. coli*, *Klebsiella*, and AIEC) ^30–33^ have evolved additional mechanisms to scavenge iron in the inflamed gut and evade lipocalin-2. These pathogens produce one or more “stealth” siderophores: salmochelins (GlcEnt), catecholate siderophores generated through the glucosylation of Ent ^34,35^, as well as the structurally distinct siderophores aerobactin (a hydroxamate-type siderophore) and yersiniabactin (a mixed-type siderophore) ^36^.

Iron deficiency is a common comorbidity in patients with IBD ^37^. These patients also exhibit an increase of *Enterobacteriaceae*, including AIEC, which harbor a diverse repertoire of siderophore synthesis and uptake machineries. These iron acquisition mechanisms likely enhance the ability of AIEC and related *Enterobacteriaceae* to colonize and persist in the iron-limited environment of the inflamed gut ^38–40^. Although a diversity of siderophore systems has been described in IBD-associated *E. coli*, the contribution of different siderophore types to enteric bacteria colonization in the inflamed gut remains undefined. Previous work demonstrated that production of GlcEnt enables the pathogen *Salmonella enterica* serovar Typhimurium (*S.* Typhimurium) to thrive in the inflamed gut and by evading lipocalin-2-mediated iron sequestration ^30^. Building on these findings, we previously designed and implemented a siderophore-based vaccine consisting of Ent conjugated to the cholera-toxin subunit B (CTB-Ent) and first demonstrated that it induced a mucosal antibody response against Ent and GlcEnt that was protective during STm infection ^41^. Importantly, in a subsequent study, CTB-Ent immunization also reduced AIEC colonization and promoted restoration of the mucosal barrier in mouse models of IBD ^42^, indicating that Ent and GlcEnt may be conserved and promising targets for the design of narrow-spectrum antimicrobials against *Enterobacteriaceae*.

Here, we show that four catecholate siderophore receptors have a redundant role in mediating AIEC iron acquisition *in vitro* and promote AIEC colonization of the inflamed gut. These results are consistent with the protective role of CTB-Ent immunization in our mouse models of colitis and AIEC infection. By performing shotgun metagenomics on samples from the mucosal surface and the lumen of the colon, we provide evidence that the protective effect of CTB-Ent immunization is most potent and selective against mucosal AIEC colonization, causing only minimal changes in gut microbiome composition. Finally, based on our metagenomics prediction, we identified *Lactobacillaceae* as a commensal microbe that increased during CTB-Ent immunization, and found that administration of representative member *Limosilactobacillus reuteri* to gnotobiotic *Il10*^-/-^ mice reduced AIEC colonization and mucosal association. Conversely, *Akkermansia muciniphila* increased in mock-immunized mice during AIEC-induced colitis, and its administration enhanced AIEC pathogenesis in gnotobiotic mice.

Together, our findings indicate that CTB-Ent immunization is a promising strategy for selective targeting of AIEC and likely other *Enterobacteriaceae* that rely on highly conserved catecholate siderophore systems to acquire iron in the inflamed gut, thereby enabling precision editing of the gut microbiota during colitis.

## RESULTS

### Catecholate siderophore uptake provides a competitive advantage to AIEC during colitis

The genomes of many pathogenic *Enterobacteriaceae*, including AIEC, encode multiple iron uptake systems that enable acquisition when this metal nutrient is limited. However, the importance of these iron uptake systems to AIEC pathogenesis in the gut remains unclear. The model AIEC strain NRG857c encodes a diverse set of genes for the synthesis and uptake of siderophores and heme ^33^. To test the contribution of these different iron acquisition systems to AIEC colonization, we generated several mutants with genomic deletions of outer membrane receptors for catecholate siderophores, non-catecholate siderophores, and heme (**Table S1**). Specifically, we deleted the four catecholate siderophore receptors FepA and IroN (the primary receptors for Ent and GlcEnt), and CirA and Fiu (the receptors for the Ent breakdown product dihydroxybenzoylserine); the yersiniabactin receptor FyuA; the aerobactin receptor IutA for aerobactin; and the ChuA receptor for heme (see **Fig. S1A**).

We first compared the growth of several AIEC iron uptake mutants to wild-type (WT) under iron-replete (nutrient broth) and iron-limited (nutrient broth supplemented with the iron chelator 2,2’-dipyridyl) conditions (**Fig. S1B-C**). In iron-replete conditions, all mutants grew to levels that were generally comparable to wild-type, except for a minor growth defect for the *fepA iroN cirA fiu* quadruple mutant (lacking the four catecholate siderophore receptors) and for the *fyuA chuA iutA* triple mutant (lacking the receptors for yersiniabactin, heme, and aerobactin). In iron-limited conditions, only the *fepA iroN cirA fiu* quadruple mutant exhibited a marked growth defect. The *fepA iroN* double mutant, and the *fyuA chuA iutA* triple mutant exhibited a mild-to-moderate growth defect, and only the *fepA iroN cirA fiu* quadruple mutant exhibited a marked growth defect (**Fig. S1B,C**). Thus, the ability to acquire iron via the four catecholate siderophore receptors appears to be particularly important for AIEC growth in an iron-limited environment. To further investigate whether these four catecholate receptors have a redundant role in Ent acquisition, we examined Ent transport in AIEC with the stable iron isotope ^57^Fe. When AIEC WT or single and double catecholate siderophore receptor mutants were grown in modified M9 minimal media supplemented with iron-bound Ent (^57^Fe-Ent), cell-associated ^57^Fe was detected (**Fig. S1D**). Only the *fepA iroN cirA fiu* mutant was unable to import ^57^Fe-Ent, further indicating that these four receptors play a redundant role for catecholate siderophore uptake in AIEC (**Fig. S1D**).

Next, we tested whether iron acquisition via the four catecholate siderophore receptors promotes AIEC colonization during colitis, when iron is limited ^30,43^. To this end, we used *Il10*^-/-^ mice, which lack the anti-inflammatory cytokine IL-10 and serve as a well-established genetic model of IBD ^44^. In our facility, *Il10*^-/-^ mice are not colonized with *Enterobacteriaceae* or *Helicobacteraceae* and do not spontaneously develop signs of disease. To accelerate the onset of colitis, we fed *Il10*^-/-^ mice a diet containing the nonsteroidal anti-inflammatory drug piroxicam for 10 days before AIEC infection ^42,45,46^. Groups of mice were then orally infected with a 1:1 mixture (10^9^ colony-forming units (CFU) total) of AIEC WT and one of the following strains: a *fepA iroN* mutant, which lacks the primary receptors for Ent and GlcEnt; a *cirA fiu* mutant, which lacks the receptors for the Ent breakdown product dihydroxybenzoylserine ^47^; a *fepA iroN cirA fiu* quadruple mutant, lacking the four catecholate siderophore receptors; or a triple mutant in the receptors for yersiniabactin, heme, and aerobactin (*fyuA chuA iutA*) (**Fig. 1A,B** and **S1A**). At days 1-5 post-infection, we observed comparable fecal colonization levels of AIEC WT and each of the catecholate siderophore double mutants (*fepA iroN* or *cirA fiu*; **Fig. 1C,D** and **S2A,B**). The *fyuA chuA iutA* triple mutant in the non-catecholate siderophore receptors (*fyuA iutA*) and the heme receptor (*chuA*) was significantly attenuated in competition with WT, although not in all mice and mostly at later timepoints (**Fig. 1F** and **S2D**). In contrast, the *fepA iroN cirA fiu* quadruple mutant, which lacks all four catecholate siderophore uptake receptors, was significantly and consistently attenuated in all mice at all timepoints (**Fig. 1E** and **S2C**). Taken together, these results demonstrate that iron acquisition, particularly via catecholate siderophores, is an important mechanism for AIEC colonization of the inflamed gut. Moreover, our findings reveal a previously unappreciated functional redundancy *in vivo* among the four catecholate siderophore receptors FepA, IroN, CirA, and Fiu.

**Figure 1.**
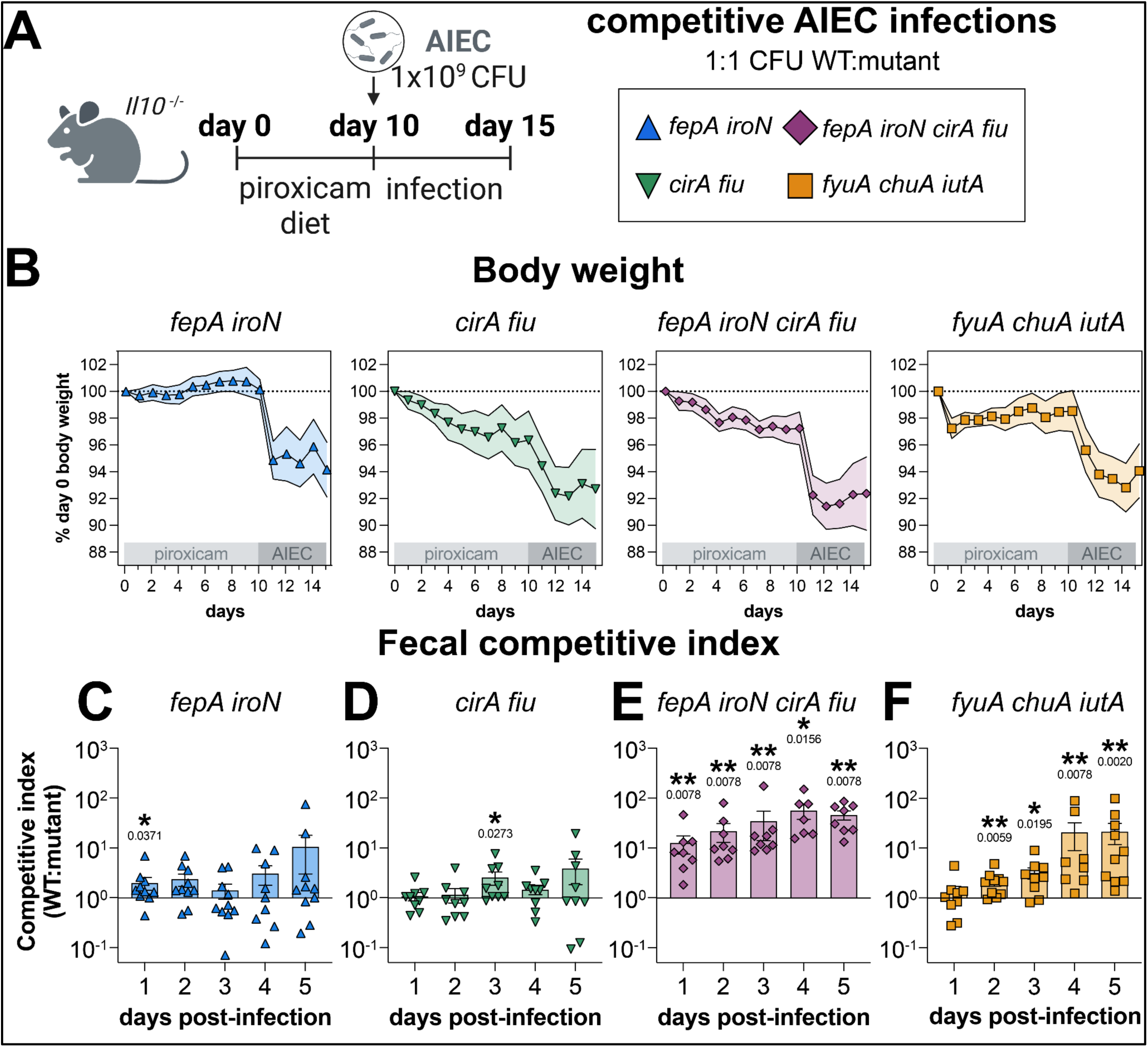
Catecholate siderophore receptors promote AIEC colonization during colitis. (**A**) Schematic representation of experimental design for AIEC competitive infections in the *Il10*^-/-^ piroxicam-accelerated colitis model. (**B**) Body weight following piroxicam administration and AIEC infection. Points represent mean percent day 0 body weight and shaded regions show SEM. (**C-F**) Fecal competitive indexes (WT:mutant AIEC CFU/mg) in mice co-infected with AIEC WT and (**C**) AIEC *fepA iroN,* (**D**) AIEC *cirA fiu,* (**E**) AIEC *fepA iroN cirA fiu,* or (**F**) AIEC *fyuA chuA iutA*. Bars show mean, points represent CFU measurements for individual animals, and lines connect paired measurements, statistical comparisons made by one-sample multiple Wilcoxon tests.

### Immunization against Ent confers protection against luminal and mucosal AIEC colonization while maintaining gut microbiota composition

We previously developed a catecholate siderophore-based vaccine by linking the non-immunogenic Ent to the carrier protein, cholera toxin B subunit (CTB-Ent) ^48^. Intranasal immunization with CTB-Ent triggered the production of antibodies that recognize Ent and GlcEnt and conferred protection against infection with *S.* Typhimurium and AIEC ^41,42^. As Ent and GlcEnt are produced only by *Enterobacteriaceae* in the gut, we previously postulated that CTB-Ent immunization more selectively targets these organisms, while leaving the commensal microbiota largely intact ^48^. Moreover, fluorescence *in situ* hybridization (FISH) of colon tissue showed that AIEC was preferentially excluded from the colonic mucosal surface in CTB-Ent-immunized mice as compared to CTB controls ^42^, suggesting that immunization may drive distinct microbiota changes in the colonic lumen versus the mucosal surface. In this study, we thus sought to elucidate the impact of CTB-Ent immunization on AIEC spatial localization and on the gut microbiota, by comparing fecal samples and mucosal scrapings.

Using CTB-Ent or CTB (control), we immunized groups of *Il10*^-/-^ mice intranasally twice, two weeks apart, as previously described ^41,42^. Two weeks after the second dose, mice were fed a piroxicam diet to accelerate colitis for 10 days, then infected with AIEC and switched to regular chow for 5 days (**Fig. 2A**). In line with our prior study ^49^, mice immunized with CTB-Ent exhibited reduced weight loss, lower fecal AIEC colonization over time compared to mice that were mock-immunized with CTB alone, and reduced intestinal inflammation at endpoint (**Fig. 2B-C** and **Fig. S3**). At day 5 p.i., mice that were immunized with CTB-Ent had significantly lower AIEC CFUs in feces, colon mucosal scrapings, and colon tissue samples as compared to the CTB control. As predicted by our FISH analysis ^42^, the difference in AIEC colonization between CTB-Ent and CTB control was most pronounced in mucosal scrapings and colon tissue samples (**Fig. 2D-E**).

**Figure 2.**
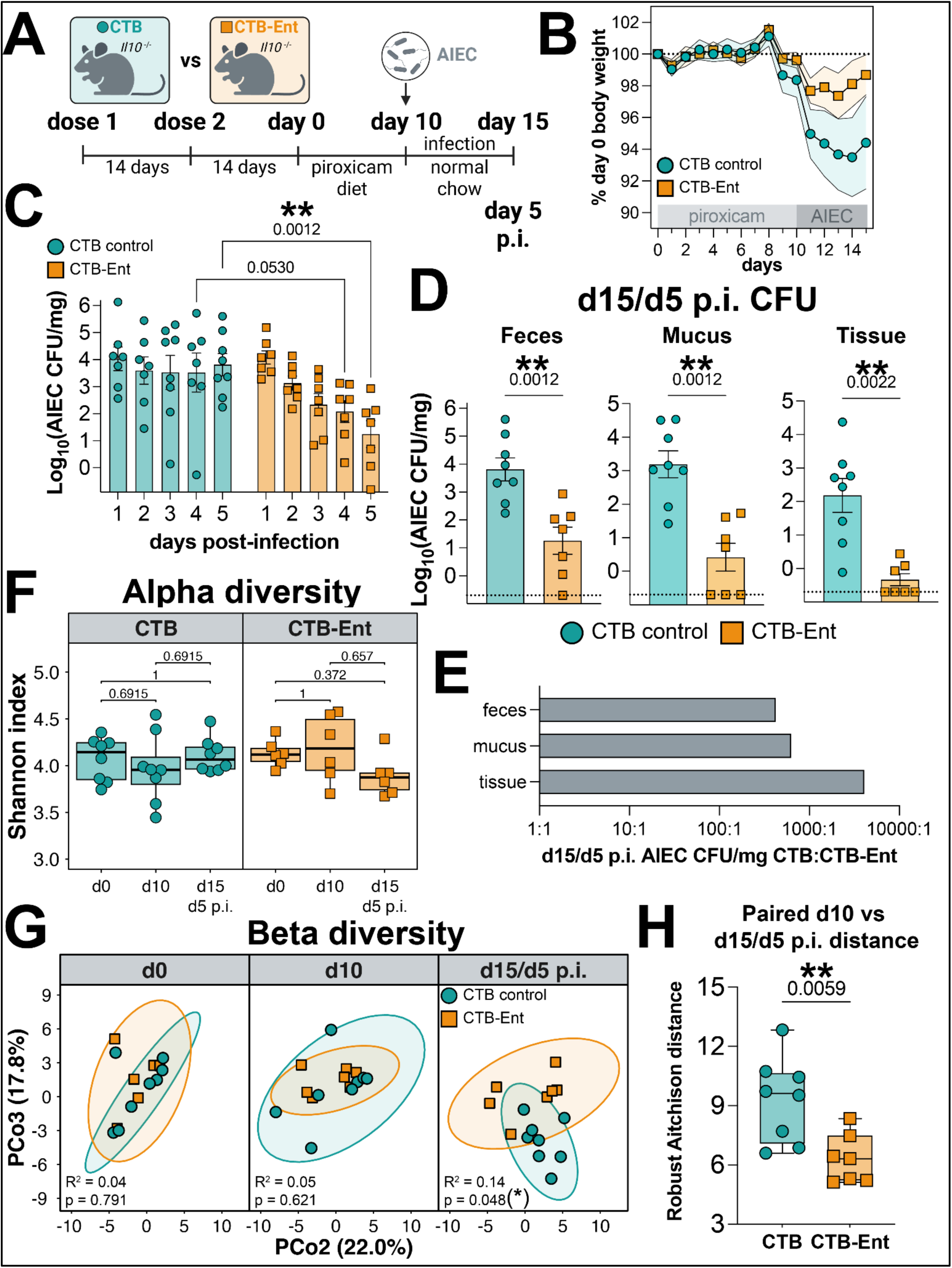
CTB-Ent immunization protects from AIEC colonization and dysbiosis. (**A**) Schematic representing CTB-Ent immunization and AIEC infection in the *Il10*^-/-^ piroxicam-accelerated colitis model. (**B**) Body weight across piroxicam administration and AIEC infection. Points represent the mean percent of day 0 body weight, and shaded regions show SEM. (**C**) Log_10_ fecal AIEC CFU/mg in CTB-Ent compared to CTB control. (**D**) Log_10_ day 5 p.i. AIEC CFU/mg between CTB-Ent and CTB control (left) feces, (middle) colon mucosal scrapings, and (right) colon tissue, and (E) mean ratio of CTB to CTB-Ent CFU/mg for each sample type at day 5 p.i. AIEC. (**F-H**) Microbiome diversity from fecal shotgun metagenomic profiles in CTB vs CTB-Ent at days 0, 10, and 15 (day 5 p.i.). (**F**) Shannon index of fecal microbiota at d0, d10, and d15. Points represent paired samples collected from the same animal at each timepoint. (**G**) Principal coordinate analysis (PCoA) of genus-level robust Aitchison distances. Points represent individual fecal samples. Statistical comparisons made by PERMANOVA (**H**) Genus-level robust Aitchison distances pre vs post-infection (d10-d15). (**C-F**) Statistical comparisons made by (D: standard; C,F: multiple) Wilcoxon tests.

From the same experimental groups (**Fig. 2A**), we performed shotgun metagenomics sequencing on fecal samples collected at day 0 (start of piroxicam diet), day 10 (end of piroxicam diet; right before AIEC infection), and day 15 (day 5 post-AIEC infection), as well as on colonic mucosa scrapings at day 15 (experiment endpoint) (**Fig. 3A**). Analysis of microbial community diversity from longitudinal fecal sampling revealed that alpha diversity remained relatively stable across piroxicam administration and AIEC infection, independent of the immunization status (**Fig. 2F**). Similarly, longitudinal analysis of within-subject robust Aitchison distance (beta diversity) demonstrated that the microbiome of mice immunized with either CTB (control) or CTB-Ent exhibited comparable changes during piroxicam treatment (**Fig. 2G**). These results are in agreement with a previous report showing that piroxicam administration did not affect the overall microbiome structure ^50^.

**Figure 3.**
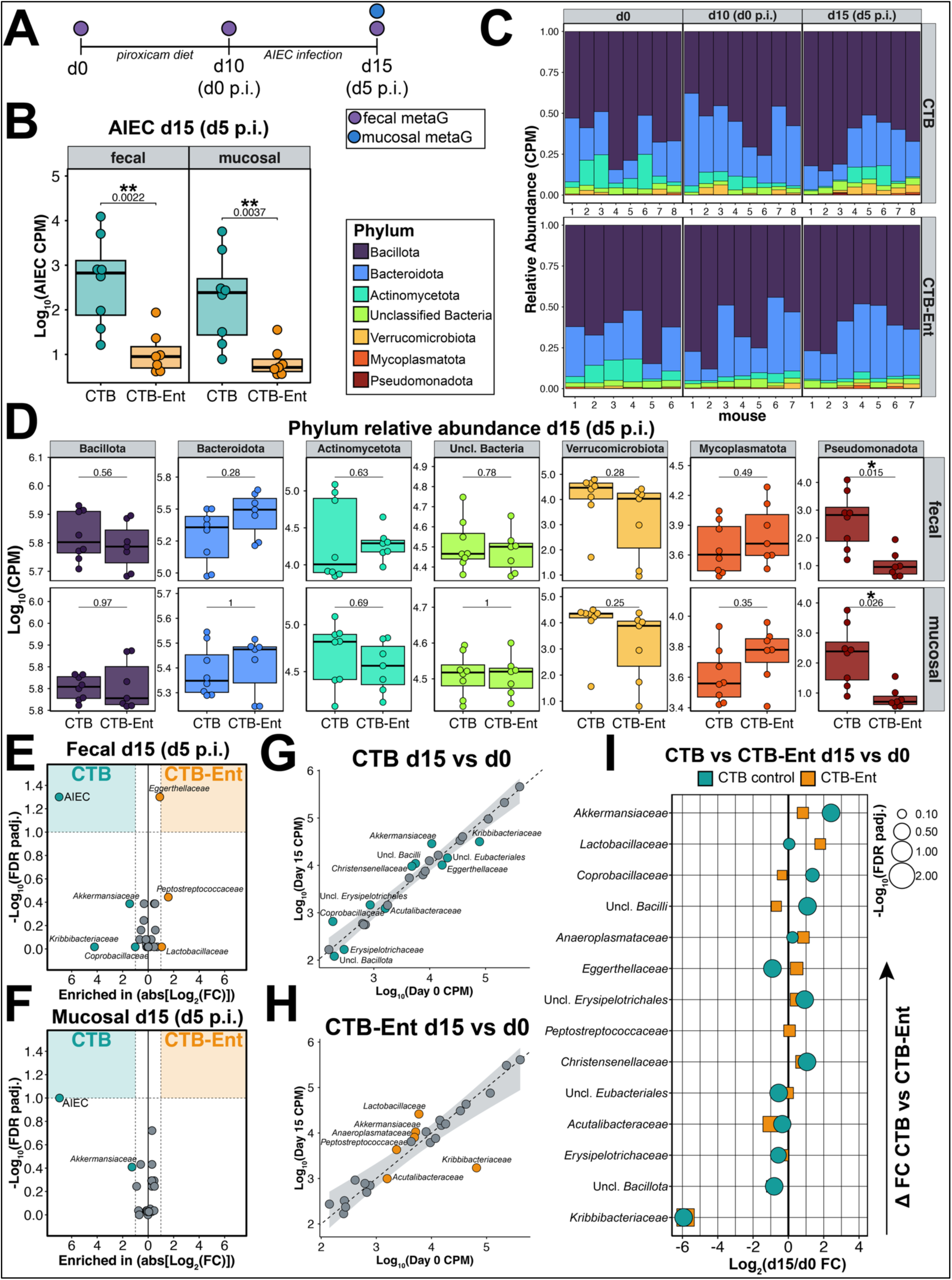
CTB-Ent gut microbiota is specifically depleted in AIEC and is distinguished by minor shifts in gut commensals. (**A**) Shotgun metagenomics analysis from fecal samples collected on days 0, 10, and 15 and colon mucosal scrapings collected from CTB control and CTB-Ent immunized mice. Raw count data normalized by counts per million (CPM). (**B**) Boxplots of day 15 (day 5 p.i.) AIEC relative abundance in feces (left) and colon mucus (right) in CTB vs CTB-Ent. (**C**) Taxa barplot of phylum-level fecal microbiota composition between CTB (top) and CTB-Ent (bottom) at (left to right) days 0, 10 and 15 of experiment. (**D**) Boxplots of phylum-level relative abundance for day 15 (day 5 p.i.) fecal (top) and mucosal (bottom) microbiota compared between CTB and CTB-Ent. (**E-F**) Volcano plot of day 15 (day 5 p.i.) family-level (**E**) fecal and (**F**) mucosal microbiota profiles between CTB (left) and CTB-Ent (right). (x-axis) log_2_-transformed fold change of mean CTB-Ent/CTB per-family relative abundances displayed as absolute values (increasing to the left is enriched in CTB; increasing to the right is enriched in CTB-Ent). (y-axis) negative log_10_-transformed FDR-adjusted p-values from per-family Wilcoxon rank-sum tests between CTB and CTB-Ent. Dashed lines and shaded regions represent thresholds for statistical significance: log_2_(FC) > |1| and FDR-adjusted p < 0.1. (**G-H**) Linear regressions of day 15 family-level relative abundances excluding AIEC between (x-axis) day 0 and (y-axis) day 15 for (**G**) CTB and (**H**) CTB-Ent. Points represent mean relative abundance and the shaded region represents the 99% confidence interval. Points within the confidence interval are colored in grey. (**I**) Bubble plot of family-level fold change in relative abundance between day 0 and day 15. (x-axis) Median log_2_-normalized within-group fold-change between day 0 and day 15 for CTB and CTB-Ent. Points represent individual genera and point size represents FDR-adjusted p-value from Wilcoxon rank-sum tests of day 0 vs day 15 relative abundances for CTB and CTB-Ent, respectively.

At the taxonomic level, minor differences were detected between day 0 and day 10 after piroxicam (**Fig. S5B**), with small but significant increases observed in the families *Peptostreptococcaceae* and *Christensenellaceae* and decreases in *Turicibacteraceae* and *Kribbibacteraceae* in both CTB and CTB-Ent immunized mice at day 10 (**Fig. S5C**). While taxonomic composition was minimally affected by piroxicam, application of functional annotation to sequence data revealed piroxicam-associated shifts in gene ontology (GO) terms between day 0 and day 10 (**Fig. S5A**). In particular, before piroxicam treatment, the fecal microbiome was enriched for GO terms associated with the biosynthesis of putrescine (**Fig. S5A**), a polyamine with important roles in suppressing inflammatory responses, that is depleted during colitis ^51^. In contrast, 10 days after piroxicam treatment, the microbiome showed an increase in GO terms associated with oxidative stress (*e.g.*, cellular oxidant detoxification, UV protection) (**Fig. S5A**). These pathway-level shifts were associated with modest changes at the family level (**Fig. S5B**), and day 10 microbiome profiles displayed small but significant enrichment in the families *Christensenellaceae* and *Peptostreptococcaceae* and a decrease in *Kribbibacteriaceae* and *Turicibacteriaceae* (**Fig. S5C**). Collectively, these pathway-level shifts suggest that piroxicam-associated inflammation creates an environment permissive to AIEC metabolism, including increased oxygen availability and access to preferred carbon sources.

After AIEC infection, the microbiome beta diversity (Aitchison distance) increased in CTB control mice, but not in CTB-Ent-immunized mice (**Fig. 2H**). This observation was further supported by principal coordinate analysis of the fecal microbiota, which revealed a distinct profile in CTB control samples at day 15 (5 days post-AIEC infection). In contrast, microbiome composition remained stable in CTB-Ent-immunized mice during infection (**Fig. 2H**). Thus, mice immunized with CTB-Ent are protected against AIEC colonization and maintain the overall gut microbiota composition.

### AIEC infection promotes the enrichment of *Akkermansiaceae* in CTB control mice, and of *Lactobacillaceae* in CTB-Ent-immunized mice

To further discern the impact of siderophore immunization on microbiota structure during AIEC infection, we compared relative abundances of bacterial taxa in fecal samples and mucosal scrapings of mice immunized with CTB-Ent in comparison to CTB controls. We noted that AIEC was the major driver of compositional difference between the CTB and CTB-Ent microbiome also on the species-level (**Fig. S4A-B**); for clarity, we conducted subsequent analyses at higher taxonomic levels (**Fig. 3**). AIEC relative abundance was significantly reduced at day 15 (day 5 p.i.) in the fecal samples and mucosal scrapings of mice immunized with CTB-Ent (**Fig. 3B**). These changes were inferred by the analysis of phylum *Pseudomonadota* (**Fig. 3D**), of which AIEC was the only member detected in the sequence data. Apart from this difference, broad phylum-level fecal microbiome composition remained stable over time in both CTB control and CTB-Ent-immunized mice (**Fig. 3C**), with the exception of a modest enrichment for the phylum *Verrucomicrobiota* in CTB-control mice, and a tendency to higher relative abundances of *Bacteroidota* and *Mycoplasmatota* in CTB-Ent-immunized mice (**Fig. 3D**). Analysis of family-level microbiome composition at day 15 (day 5 p.i.) again revealed that *Enterobacteriaceae* (single member AIEC) was the only significant differentially abundant taxon in the feces when comparing CTB control and CTB-Ent-immunized mice, with a similar trend (FDR-adjusted p = 0.1) in mucosal scrapings (**Fig. 3E, F**). These results indicate that, in a colitis model, CTB-Ent immunization more selectively targets *Enterobacteriaceae* within a complex gut microbial community.

We also observed that some bacterial families showed an association with disease severity (measured as weight loss) after 10 days of piroxicam treatment, before AIEC infection (**Fig. S6**). Among bacterial families that varied during piroxicam administration (**Fig. S5B**), abundances of *Akkermansiaceae* and *Christensenellaceae* were significantly and negatively associated with body weight change during piroxicam treatment (**Fig. S6C**). Importantly, these piroxicam-associated microbiome changes were shared between CTB and CTB-Ent immunized groups (**Fig. 2G,S6A**), and the overall shift in community composition during piroxicam administration was not associated with weight loss or AIEC engraftment (**Fig. S6B**). This observation suggests that members of the gut microbiota may independently contribute to disease severity, potentially exacerbating AIEC-associated pathology.

Although AIEC relative abundance was the only significant difference in the gut microbiome between CTB control and CTB-Ent-immunized mice, there were some temporal changes in a few other members of the gut microbiota within each group. A comparison of family-level relative abundance between day 0 (before piroxicam treatment) and day 15 (day 5 p.i.) within CTB and CTB-Ent-immunized groups revealed an increase in *Akkermansiaceae* and a decrease in *Kribbibacteriaceae* in both groups (**Fig. 3G-H**). In CTB-control-immunized mice, the day 15 (day 5 p.i.) fecal microbiome was also enriched for unclassified *Bacilli*, *Christensenellaceae*, *Coprobacillaceae*, and unclassified *Erysipelotrichales* (**Fig. 3G**). In contrast, mice immunized with CTB-Ent were uniquely enriched for *Lactobacillaceae* at the same timepoint (**Fig. 3H**). Ranked by difference in fold change between CTB controls and CTB-Ent-immunized mice, the family *Akkermansiaceae* was most enriched in CTB control groups, and *Lactobacillaceae* in CTB-Ent groups, at day 5 post-AIEC infection (**Fig. 3I**).

To test whether the observed shifts in the commensal microbiota contributed to the difference in disease severity between the CTB-immunized and the CTB-Ent-immunized mice, we selected two representative commensal bacteria: *Akkermansia muciniphila* (*Akkermansiaceae* enriched in CTB controls), and *Limosilactobacillus reuteri* (*Lactobacillaceae* enriched in the CTB-Ent group). We then colonized germ-free (GF) *Il10^-/-^* mice with AIEC, *A. muciniphila*, or *L. reuteri* alone, or with AIEC plus *A. muciniphila*, or AIEC plus *L. reuteri* (**Fig. 4A**). After 4 days, *Il10^-/-^* mice monocolonized with either AIEC or *A. muciniphila* lost significantly more weight compared to the *L. reuteri* monocolonized group, and this trend continued throughout the 7-day experimental interval (**Fig. 4B**). Remarkably, *Il10^-/-^* mice co-colonized with AIEC and *A. muciniphila* exhibited greater weight loss relative to AIEC monocolonized mice, whereas co-colonization with *L. reuteri* ameliorated AIEC-associated weight loss (**Fig. 4C**). *A. muciniphila* co-colonization also significantly enhanced AIEC fecal burdens by day 7 post-colonization relative to AIEC monocolonization and co-colonization with *L. reuteri* (**Fig. 4D**). In agreement with these results, host inflammatory markers measured by qPCR revealed similar levels of expression in AIEC monocolonized and AIEC + *A. muciniphila* co-colonized mice (**Fig. 4E**). In contrast, mice co-colonized with AIEC + *L. reuteri* exhibited significantly lower markers of neutrophilic inflammation, as evidenced by diminished expression of inducible nitric oxide synthase 2 (*Nos2*), the neutrophil marker *Ly6g*, and the neutrophil antimicrobial protein calprotectin (*S100a9*), whereas the antimicrobial protein lipocalin-2 (*Lcn2*) did not change significantly (**Fig. 4E**).

**Figure 4.**
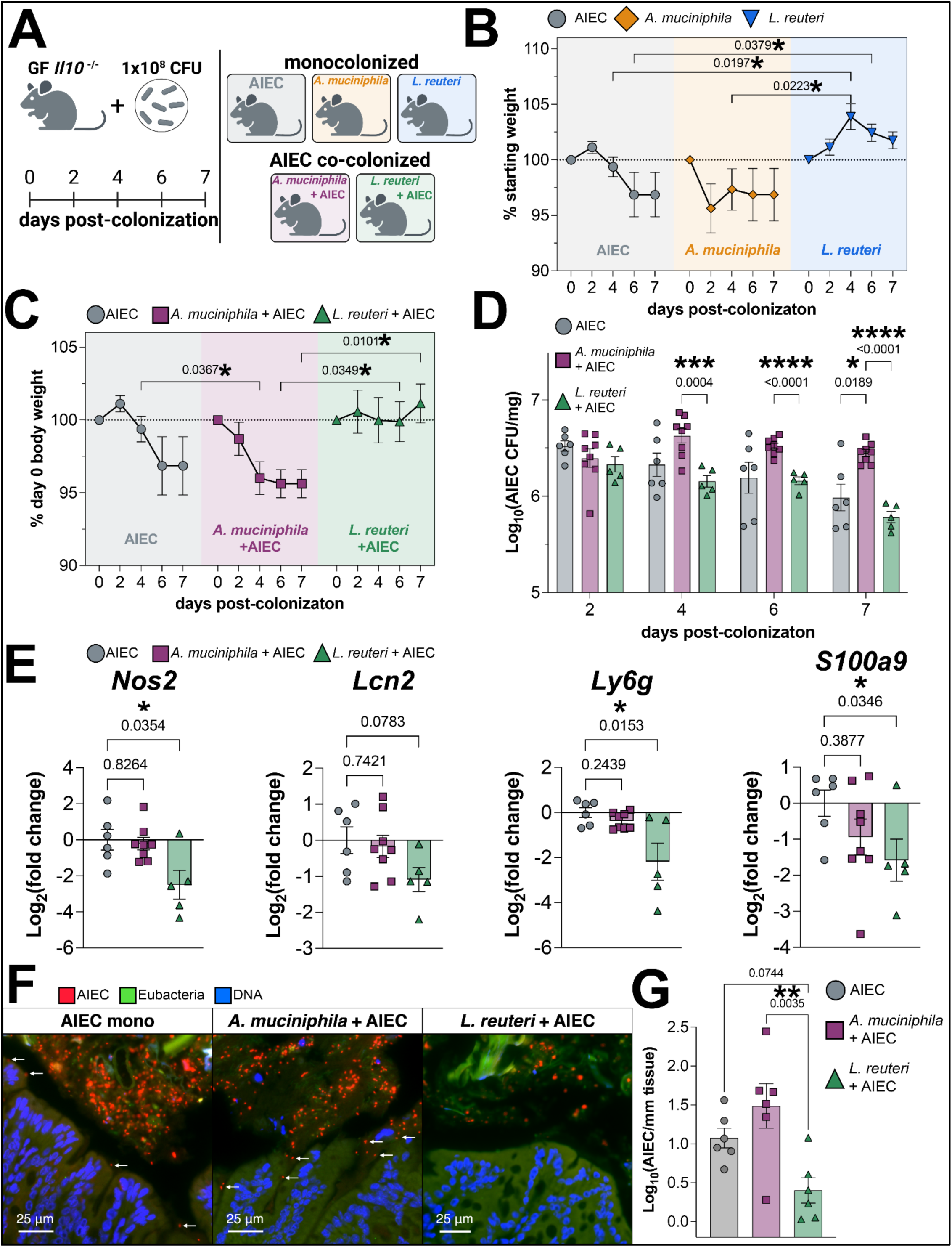
Differential contributions of CTB and CTB-Ent-associated microbiota shifts to AIEC pathogenesis in gnotobiotic *Il10^-/-^* mice. (**A**) Schematic of experiment design. Germ-free *Il10^-/-^* mice were monocolonized with 10^8^ CFU of AIEC, *Akkermansia muciniphila*, or *Limosilactobacillus reuteri,* or co-colonized with a 1:1 mix of 2×10^8^ CFU total AIEC + *Akkermansia* (model CTB control microbiota) or AIEC + *Limosilactobacillus reuteri* (model CTB-Ent microbiota). After inoculation, body weight and fecal AIEC CFU were measured at days 2, 4, 6 and 7 p.i. and fecal samples for metagenomics were collected at days 4 and 7 p.i. (**B,C**) Mouse body weight as a percentage of day 0 for (**B**) monocolonized and (**C**) AIEC co-colonized animals. Weights compared to AIEC monocolonized animals by ANOVA. (**D**) Log_10_-normalized fecal AIEC CFU/mg between AIEC monocolonized and co-colonized animals compared by ANOVA. (**E**) Relative expression of host genes in distal colon samples from AIEC monocolonized and co-colonized animals quantified by qPCR and represented as Log_2_-normalized fold change relative to mean AIEC monocolonized expression. Fold change values compared to AIEC monocolonized animals by Kruskal-Wallis tests. (**F**) Representative images from fluorescence *in situ* hybridization (FISH) staining of distal colon sections from Germ-free *Il10^-/-^* mice monocolonized with (left panel) AIEC, or co-colonized with (middle panel) AIEC + *A. muciniphila* or (right panel) AIEC + *L. reuteri*. (**G**) Quantification of tissue-associated AIEC from n=3 regions of FISH images in n=2 independent samples.

As AIEC is thought to exacerbate inflammation by its close association with the intestinal mucosa, we sought to investigate whether co-colonization with *A. muciniphila* or *L. reuteri* impacted the spatial localization of AIEC within the gut. We performed fluorescence *in situ* hybridization (FISH) of distal colon sections, using a probe that detects AIEC ^42^ (**Fig. 4F**). Tissue-associated AIEC was higher in *Il10^-/-^* mice co-colonized with AIEC and *A. muciniphila*, but it was reduced in mice co-colonized with AIEC and *L. reuteri*, relative to mice monocolonized with AIEC (**Fig. 4F,G**). Collectively, these data suggest that microbiota shifts associated with CTB (control) immunization (e.g., an increase in *A. muciniphila*) contribute to disease severity during AIEC infection. In contrast, microbiota changes in mice immunized with CTB-Ent (e.g., the increase in *L. reuteri*) may be protective.

## DISCUSSION

Microbiota-sparing therapies that target the microbial component of disease pathogenesis are an area of intense interest in IBD ^52^. The therapeutic role of traditional antibiotics has been limited by their modest clinical benefits and their tendency to further disrupt an already imbalanced gut microbiome ^53^. Therefore, narrow-spectrum antimicrobial strategies are needed to selectively target the microbes that contribute most to disease severity. In this regard, AIEC presents an attractive therapeutic target, as this pathovar is present in up to 60% of Crohn’s disease patients and its abundance is known to fluctuate along with the relapsing and remitting pattern of IBD symptoms ^18,20^. Here, we demonstrate that four catecholate siderophore receptors promote AIEC growth in the inflamed gut, and that targeting catecholate siderophores through immunization selectively reduces AIEC colonization within a complex endogenous gut microbial community.

Immunization with CTB-Ent confers protection during infection with *S.* Typhimurium or AIEC ^41,42^. It is intriguing that this strategy is successful in the context of AIEC infection, given the high diversity of iron acquisition systems encoded within the AIEC genome ^14,33^ in comparison, for instance, to *S.* Typhimurium^30^. Indeed, AIEC and other IBD-associated *E. coli* produce several outer membrane transporters for the uptake of siderophores and heme ^54^. Of note, the AIEC strain used in this study produces at least eight independent receptors for iron acquisition, including four dedicated to catecholate siderophores and an additional four dedicated to the stealth siderophores yersiniabactin, aerobactin, ferrichrome, and the iron-chelate complex heme, which together confer resistance to host lipocalin-2. Because CTB-Ent immunization targets only catecholate siderophores and we observed a reduction of AIEC colonization in the inflamed gut ^49^, we hypothesized that catecholate siderophore uptake represents the major iron acquisition pathway for AIEC in the inflamed gut. Consistent with this notion, deletion of all four catecholate siderophore uptake systems significantly attenuated AIEC colonization in competition with WT in all mice at every timepoint analyzed. Unexpectedly, deletion of the two catecholate siderophore receptors FepA and IroN did not significantly impact AIEC colonization, which led us to investigate a potential role for Cir and Fiu. We found that these two receptors contribute to enterobactin uptake *in vitro*, and promote AIEC growth *in vivo*. Of note, we are not aware of other studies in which Cir and Fiu were found to contribute to iron acquisition *in vivo*. Together, our results suggest that the catecholates are not only the most physiologically important siderophore class for AIEC in the inflamed gut, but also imply that these four receptors produced by AIEC contribute redundantly to catecholate siderophore-mediated iron uptake *in vivo*. These findings further suggest that the protection mediated by CTB-Ent immunization likely results from blockade of iron acquisition across all catecholate siderophore uptake systems. This is in contrast, for instance, to the siderophore antibiotic cefiderocol, which preferentially targets Cir-mediated uptake, creating selective pressure for mutations in this receptor ^55,56^. We also found that an AIEC mutant in three non-catecholate iron acquisition systems was comparatively less attenuated in our model *in vivo*. Nevertheless, future immunization strategies could consider targeting Ent/GlcEnt together with these additional systems. A prior study with uropathogenic *E. coli* found that yersiniabactin and aerobactin can also be used for immunization ^57^, showing the promise of this approach.

The success of narrow-spectrum antimicrobials relies on their selectivity for the intended target organism. Beyond investigating the efficacy of CTB-Ent immunization, we sought to determine whether it was selective for AIEC during colitis. As many bacterial and fungal pathogens leverage siderophore-mediated iron uptake as virulence factors, siderophores have emerged as attractive therapeutic targets ^58^. The promise of siderophore-based therapies as antimicrobials is further enhanced in the context of class IIb microcins, endogenous antimicrobial peptide-siderophore conjugates that aid in competition between *Enterobacteriaceae* ^59^. However, while often proposed as a precision antimicrobial strategy, the specificity of siderophore-based therapies in the context of complex *in vivo* microbial communities has yet to undergo robust evaluation. Indeed, although Ent is uniquely produced by *Enterobacteriaceae*, recent studies have revealed that Ent can be used by unrelated microbes as a xenosiderophore, which might complicate the paradigm of siderophore-based therapies as selective antimicrobials ^60–62^. Therefore, we sought to perform, to our knowledge, the most comprehensive evaluation of the specificity of a siderophore-based antimicrobial to date by applying longitudinal fecal shotgun metagenomic profiling from mice immunized with CTB-Ent or mock-immunized with CTB. The result of this analysis first revealed no baseline differences between the microbiome of CTB-immunized control mice and that of CTB-Ent-immunized littermate mice, indicating that CTB-Ent immunization does not affect the microbiome during homeostasis. Furthermore, after infection, AIEC was the only significant differentially abundant organism between the CTB- and CTB-Ent immunized groups, indicating the selectivity of this immunization strategy against its target organism. These results provide a strong proof-of-principle that CTB-Ent immunization can be used to antagonize a single organism, AIEC, within an endogenous microbiota *in vivo*.

Beyond its therapeutic applications, the specific impact of CTB-Ent immunization against AIEC affords a unique opportunity to study the dynamics of the microbiota while just a single organism is antagonized. In contrast to broad-spectrum antimicrobials that act as a global perturbation by targeting many members of a microbial community, our metagenomics data indicate that CTB-Ent acts as a precision perturbation by exerting stress on AIEC alone. Therefore, CTB-Ent immunization may cause a unique ecological pressure associated with distinct collateral effects on the surrounding commensal microbiota. Although AIEC was the only organism with significantly altered abundance between treatment groups at day 15, some immunization-specific alterations to the microbiota were also observed. In particular, the family *Akkermansiaceae* was uniquely enriched in CTB control mice. The sole genus within this family, *Akkermansia*, is a mucolytic genus of Gram-negative obligate anaerobes that has been reported to both ameliorate and exacerbate colitis in a context-dependent manner ^63,64^. In our study, *Akkermansia* abundance was also associated with increased weight loss during piroxicam administration, before AIEC infection. Together, these observations suggest that the increased abundance of *Akkermansia* in CTB controls may play a causal role in exacerbating AIEC-associated pathology. In contrast, the family *Lactobacillaceae* was enriched in the day 15 fecal metagenome of CTB-Ent immunized mice. This finding was consistent with our previous investigation of the protective role of CTB-Ent immunization during *S.* Typhimurium infection, in which we also detected an increase in *Lactobacillus* by 16S amplicon sequencing ^41^. Of note, lactobacilli are broadly considered beneficial bacteria and are often used as probiotics ^65^, raising the possibility that their administration could be beneficial during AIEC infection.

In agreement with the observed immunization-specific shifts in the commensal microbiota, monocolonization of germ-free *Il10^-/-^* mice with AIEC or *A. muciniphila* resulted in increased disease severity relative to mice monocolonized with *L. reuteri*. Somewhat unexpectedly, *Il10^-/-^*mice monocolonized with *A. muciniphila* lost weight more rapidly than those monocolonized with AIEC. This observation is also in line with the pre-infection association between weight loss and *A. muciniphila* abundance in our specific pathogen-free (SPF) *Il10*^-/-^ mice treated with piroxicam. Previous studies have reported an association between *A. muciniphila* and colitis in the context of a complex microbiota or during pathogen infection, citing the predilection of this organism to degrade host mucins and compromise the integrity of the mucus layer to immunogenic microbial stimuli ^64^. However, our data indicate that *A. muciniphila* alone is sufficient to induce colitis in specific contexts, as previously reported^66^. In contrast to *A. muciniphila*, *Il10*^-/-^ mice monocolonized with *L. reuteri* did not exhibit weight loss, and co-colonization with *L. reuteri* protected AIEC-infected mice from weight loss and reduced AIEC colonization and inflammation. The mechanisms of *L. reuteri*-mediated protection in our model need to be investigated and might involve both niche exclusion and production of anti-inflammatory and antimicrobial metabolites ^67,68^.

Broadly, our study provides a proof-of-principle for the precision depletion of AIEC, a pathovar considered a trigger of disease in IBD patients, by exploiting an Achilles heel in the gut: iron acquisition via catecholate siderophores. Whereas AIEC have evolved multiple means by which to acquire catecholate siderophores in the inflamed gut, a particular strength of our approach is that it simultaneously disrupts all of them by targeting their common substrate. Moreover, we have identified *Akkermansiaceae* and *Lactobacillaceae* as organisms that respectively exacerbated or ameliorated AIEC-induced colitis in genetically prone mice, implicating these microbes as possible targets for therapeutic modulation during IBD.

### Limitations of the study

Although CTB-Ent immunization specifically reduced AIEC colonization in the context of the *Il10*^-/-^ microbiota in our colony, this finding does not preclude the possibility for off-target effects in other mouse colonies with different microbial communities. We should also note that the SPF *Il10*^-/-^ mice in which the immunization studies described here were performed are intentionally maintained without endogenous *Enterobacteriaceae*. In addition, the only Ent utilization loci annotated in our shotgun metagenomic data were within the AIEC genome. We chose to intentionally target AIEC with CTB-Ent immunization because of its known association with IBD severity. However, many *Enterobacteriaceae* participate in commensal and even mutualistic interactions with the host by contributing to colonization resistance against their pathogenic counterparts ^69–71^. In addition, while Ent uptake was originally believed to be restricted to members of the order *Enterobacterales*, recent studies have challenged this paradigm by demonstrating that some unrelated taxa leverage Ent uptake as a xenosiderophore ^72,73^. In these scenarios, CTB-Ent immunization may result in antagonism of some commensal Ent utilizers, and potentially weaken the natural barrier against pathogens conferred by the gut microbiota. Future studies of siderophore-based immunization in systems that contain commensal *Enterobacteriaceae* and unrelated organisms that utilize Ent as a xenosiderophore should be prioritized to fully elucidate the potential off-target effects of CTB-Ent immunization. Therefore, while the durability of CTB-Ent-mediated immunity has not been interrogated, its potential to promote long-term antagonism of Ent utilizers raises questions surrounding the tractability of this approach. To address this limitation, we are interested in developing CTB-Ent immunization into more transient therapeutic modalities, such as monoclonal antibodies against Ent.

## Acknowledgements

We would like to thank Dr. Brian Coombes (MacMaster University) for providing us with plasmids, tools, and protocols for making mutants in AIEC. Confocal imaging and slide scanning were done at the UCSD School of Medicine Microscopy Core, which is supported by a NINDS P30 grant (NS047101). Histology was performed at the La Jolla Institute for Immunology core and was partly supported by NIDDK Grant P30 DK120515. Sequencing was performed at the UCSD IGM Genomics Center and was partly supported by NIDDK Grant P30 DK120515.

## Funding

National Institute of Health grant F31 AI186410 (GJN) Max Kade Foundation (RRG)

Crohn’s and Colitis Foundation grant 649744 (RRG)

National Institute of Health grant T32AI007036 (GTW)

National Institute of Health grant R01AI176390 (MR and EMN)

National Institute of Health grant R37AI126277 (MR)

National Institute of Health grant R21AI186034 (MR and KZ)

National Institute of Health grant R01AI167860 (HC)

National Institute of Health grant R01AI188710 (HC)

AMED grant JP233fa627004 (MR and HC)

Chiba University-University of California-San Diego (UCSD)

Center for Mucosal Immunology, Allergy, and Vaccines (MR and HC)

## Author contributions

Conceptualization: GJN, RRG, EMN, MR Methodology: GJN, RRG, CG, GK, HC, KZ, EMN, MR

Software: GJN, KZ

Investigation: GJN, AMD, RRG, CG, VC, FAA, CJB, GK, JD, GTW, C-YH, SB

Resources: HC, KZ, EMN, MR

Visualization: GJN

Formal analysis: GJN Funding acquisition: EMN MR

Project administration: MR

Supervision: GJN, RRG, S-PN, HC, KZ, EMN, MR

Writing – original draft: GJN, MR

Writing – review & editing: GJN, AMD, RRG, CG, VC, FAA, CJB, GK, JD, GTW, C-YH, SB, S-PN, HC, KZ, EMN, MR

## Data and materials availability

All data are available in the main text or in the supplementary materials.

## MATERIALS AND METHODS

### Bacterial strains and culture conditions

Adherent-invasive *Escherichia coli* (AIEC) strain NRG857c was kindly provided by Dr. Alfredo Torres (UTMB Health) and Dr. Brian Coombes (McMaster University) ^33^. Unless otherwise indicated, AIEC wild-type and mutants were cultured overnight under aerobic conditions at 37 °C in Miller Lysogeny broth (LB) or on LB agar plates. Where indicated, media was supplemented with 30 μg/mL chloramphenicol (Cm), 10 or 15 μg/mL gentamicin (Gm_10_; Gm_15_), or 100 μg/mL kanamycin (Km). When relevant, the wild-type AIEC strain is denoted as AIEC WT. All strains are listed in **Supplemental Table 1**.

*Akkermansia muciniphila* strain BAA-835 was kindly provided by Dr. Michael Fischbach (Stanford University). *A. muciniphila* was cultivated under anaerobic conditions at 37 °C in degassed BHIS broth or on BHIS agar supplemented with 1 mM L-cysteine and 1:1000 BD BBL vitamin K1-Hemin solution ^74^. *Limosilactobacillus reuteri* was isolated from the cecal contents of mice bred and maintained in UCSD facilities. *L. reuteri* was cultivated under anaerobic conditions at 37 °C in MRS broth or on MRS agar (Millipore Sigma).

### Bacterial mutant generation

AIEC genomic deletions were constructed using the lambda Red recombineering system ^75^. Mutagenesis protocols and plasmids were generously provided by Dr. Brian Coombes (McMaster University) ^76^. In brief, primers (**Supplemental Table 2**) were designed with homology to flanking 40-50 bp genomic regions directly 5’ and 3’ of the target gene for deletion. The 3’ ends of primers were designed with homology to the gentamicin-resistance (Gm^R^) cassette in plasmid pCDF_GmFrt ^76^. Amplicons were transformed into AIEC carrying the pKD46-Km plasmid construct, and mutants recovered on LB+Gm_10_ were confirmed by PCR to verify successful insertion and directionality of the Gm^R^ resistance cassette. To generate multiple genomic deletions, strains were then cured of pKD46-Km, transformed with pCP20-Km, cured of pCP20-Km, and transformed with pKD46-Km before inserting Gm^R^ cassettes at additional genomic loci.

### Bacterial *in vitro* growth assays

Cultivation of AIEC WT and AIEC mutants under iron-limited conditions was accomplished by growing strains aerobically overnight at 37 °C in Nutrient Broth supplemented with 100 μM 2’,2’-dipyridyl (NBD broth). Overnight cultures were then pelleted, washed, serially diluted, and inoculated into nutrient broth or nutrient broth supplemented with 2,2’-dipyridyl at a density of 1×10^4^ colony-forming units (CFU) per mL ^30,43^. For growth assays, cultures were then incubated aerobically, shaking at 37 °C; and growth was monitored by optical density (OD_600_) for up to 24 hours. Growth data were then modeled using R version 4.5.0 with the Growthcurver package version 0.3.1 ^77,78^. Quantification of cell-associated iron was performed as previously described ^79^, except that the bacterial cell pellets were digested by treatment with 0.1 mL concentrated nitric acid (30 min, RT). Briefly, AIEC WT or mutants were grown aerobically overnight at 30 °C in modified M9 minimal medium, then the cultures were diluted into fresh modified M9 for an additional 3-4h at 30 °C until mid-log phase. Cultures were then pelleted, resuspended in modified M9 supplemented with 10 μM ^57^Fe^3+^-Ent and incubated aerobically, shaking at 30 °C for 30 min. After incubation, optical density (OD_600_) was measured, cultures were pelleted, washed twice in fresh modified M9 followed by one additional wash in M9 + 2% EDTA, and cell pellets were harvested. Cells were then digested in concentrated nitric acid for 30 min at room temperature, and lysates were diluted in water to a final nitric acid concentration of 5%. The resulting digests were diluted with 1.9 mL of Milli-Q water and then analyzed by inductively-coupled mass spectrometry as described ^79^.

### Mice

Specific pathogen-free (SPF) C57BL/6 *Il10^-/-^* mice were originally acquired from The Jackson Laboratory, then bred and maintained at UCSD. Mice were maintained with irradiated food (Teklad diet 2920X) and autoclaved water, and fecal samples were periodically plated on MacConkey agar to verify that mice were not colonized with *Enterobacteriaceae*. The mice were kept in a 12h light/dark cycle, at a room temperature of ∼22 °C, and ∼52% humidity. All animal experiments were approved by the Institutional Animal Care and Use Committee at the University of California San Diego (protocol number S17107). Germ-free C57BL/6 *Il10^-/-^* mice were bred and housed in flexible film isolators (Class Biologically Clean) until 6-8 weeks of age, then transferred to micro-isolator cages (Allentown) and maintained with autoclaved food (Lab Diet 5010), bedding, and water prior to the colonization experiments.

### Immunization

CTB-Ent was prepared as previously described ^41,42^. Six-week-old male *Il10^-/-^* mice were immunized by intranasal administration of 100 μg of cholera toxin subunit B (CTB; Sigma) or CTB-Ent solubilized in PBS. Mice were then administered a booster dose of 100 μg of their respective treatment 14 days after the first dose ^41,42^. Piroxicam administration began 14 days after the immunization booster.

### *Il10^-/-^* piroxicam-accelerated colitis model and AIEC infection

For competitive infections between AIEC wild-type and the iron-uptake mutants, eight to ten-week-old *Il10^-/-^* male and female mice were used. For the immunization studies, we used eight to ten-week-old *Il10^-/-^* male mice, as in our prior study we found them to exhibit a more consistent inflammatory response than female mice ^42^. All mice were switched from normal chow to a Teklad 2020X-based diet containing 100 parts-per-million (PPM) piroxicam for 10 days prior to infection ^42,80^. During this interval, diet was replaced daily, and body weight was monitored. After 10 days, mice were administered 1×10^9^ AIEC CFU by oral gavage. Body weight was monitored for 5 days following infection and fecal AIEC CFUs were quantified as described below. On day 5 post-infection, mice were euthanized by CO_2_ asphyxiation and intestinal samples were collected for histopathology, analysis of gene expression changes, as well as microbiome analysis. In some groups of mice, we performed mucosal scrapings as previously described ^81^, to quantify AIEC associated with the mucosa and to perform microbiome analysis.

### Germ-free mouse colonization

Eight-week-old germ-free *Il10^-/-^* mice were monocolonized with 1×10^8^ CFU of AIEC, *A. muciniphila*, or *L. reuteri*. For co-colonization experiments mice were colonized with 2×10^8^ CFU (1×10^8^ CFU/strain) of AIEC and *A. muciniphila* or *L. reuteri*. For 7 days following infection, body weight and fecal AIEC CFU were quantified as described below. On day 7 post-colonization, mice were euthanized by CO_2_ asphyxiation and intestinal samples were collected.

### Quantification of bacterial colonization

Prior to AIEC infection, fecal samples were collected from *Il10^-/-^*mice to screen for the absence of *Enterobacteriaceae* by plating fecal homogenates on MacConkey agar. Similarly, prior to colonization, germ-free animals were screened for cultivable bacterial colonization by plating fecal homogenates on BHIS agar. During infection/colonization, fecal AIEC CFUs were quantified by weighing and homogenizing mouse fecal pellets in 1 mL of sterile PBS. Fecal homogenates were then serially diluted with sterile PBS and plated onto LB+Cm for AIEC single infections, or LB+Cm and LB+Gm_15_ for AIEC competitive infections, then incubated at 37 °C overnight.

### Extraction of bacterial DNA from mouse intestinal samples

Fecal pellets and mucosal scrapings were collected, immediately snap-frozen in liquid nitrogen, and stored at -80 °C prior to DNA extraction. Mucosal scrapings were collected by gently scraping the colon mucosal surface with a sterile, plastic inoculating loop and transferring the contents into a cryovial prefilled with DNA/RNA Shield (Zymo Research) ^82^. DNA was extracted using the ZymoBIOMICS DNA miniprep kit (Zymo Research) per the manufacturer’s instructions. DNA concentrations were measured using the Qubit dsDNA HS Assay kit and Qubit 2.0 fluorometer (Qubit; Invitrogen).

### Shotgun metagenomics library preparation and sequencing

Metagenomics libraries were prepared using the Nextera XT library preparation kit (Illumina) per the manufacturer’s instructions and amplified with IDT for Illumina xGEN NXT UDI Primers (IDT). Library quality and size were evaluated using the 4200 TapeStation (Agilent) and concentration quantified by Qubit. Libraries were sequenced on a NovaSeq X Plus with 1.5B flow cell and PE100 run configuration to a target depth of 1×10^7^ reads per library.

### Metagenomics data processing and analysis

Adapter sequences were trimmed from raw reads using TrimGalore (Cutadapt) version 0.6.6. and quality controlled for reads with Phred score ≥20 using FastQC version 0.11.9. ^83,84^. Trimmed reads were then passed to Sylph version 0.7.0 to estimate taxonomic composition. The resulting taxonomic profile consisted of n=277 genomes not identified as *Escherichia coli*, and the reference genome for AIEC NRG857c (GCF_003833075.1) was added to the final genome set of n=278 ^85^. Genomes and associated metadata were downloaded using NCBI Datasets version 17.3.0, and a custom database was constructed using bowtie2 version 2.2.3 ^86^. Trimmed reads were then aligned using bowtie2 and count tables were generated using Woltka version 0.1.5. ^87^. Downstream processing of sequence data was performed using R version 4.5.0. and RStudio ^77^.

Analysis of microbiome diversity was performed using vegan version 2.7-2 ^88^. For quantification of community alpha diversity, raw read counts were rarefied to a depth of 5×10^6^ reads per sample. For beta diversity, robust Aitchison distances were calculated from robust center-log ratio transformed count data^89^. For analysis of taxon relative abundance, raw read counts were normalized by sum-scaling (counts per million; CPM).

### RNA extraction from mouse tissues

Distal colon sections were cleared of content, immediately snap-frozen in liquid nitrogen, and stored at -80 °C prior to RNA extraction. RNA was extracted by transferring tissues to RINO Bead Lysis kit tubes (Next Advance) and homogenizing in Tri-Reagent (Molecular Research Center) on a Bullet Blender Tissue Homogenizer (Next Advance). Lysates were then transferred to EconoSpin RNA Mini Spin Columns (Epoch Life Science Inc.), DNA was degraded using the PureLink on-column DNase Set (Life Technologies), then cDNA was synthesized from 1 μg input RNA using LunaScript RT SuperMix (NEB).

### Quantification of mouse transcripts by qPCR

Gene expression was quantified by qPCR using the PowerUp SYBR Green Master Mix (Life Technologies) and reactions were amplified on a QuantStudio 5 Real-Time PCR System (Thermo Fisher). Expression of target genes *Nos2, Lcn2*, *Ly6g*, and *S100a9* were normalized to *Actb* (β-actin). Fold change in gene expression was compared to mean expression values of AIEC monocolonized mice using the ΔΔCt method.

### Histopathology

Distal colon sections were fixed in 10% buffered formalin for 24 hours and stored in 70% ethanol prior to processing. Tissues were then embedded in paraffin according to standard procedures, sectioned at 5 μm, and stained with hematoxylin and eosin (H&E). Slides were scanned using a Axio Scan.Z1 (Zeiss) and visualized using QuPath. Inflammation was scored in a blinded fashion using a semiquantitative scoring system as described previously (maximum score = 48) ^42,90^.

### Fluorescence *in situ* hybridization

Distal colon sections were fixed in Carnoy’s fixative for 24 hours and stored in 70% ethanol prior to processing ^42,91^. Tissues were then embedded in paraffin according to standard procedures and sectioned at 5 μm. Sections were then processed and hybridized to fluorescent pan-bacterial or *Enterobacteriaceae*-specific probes as previously described ^42,91^. Fluorescence microscopy and imaging were conducted using an Olympus VS200 slide scanner. Image analysis was performed using QuPath version 0.6.0 ^92^.

### Statistical analysis

Statistical analysis was performed with GraphPad Prism 11 and R version 4.5.0. *In vitro* growth assay data were compared using ANOVA with Dunnett correction. CFU data from *in vivo* infection models were Log_10_-transformed and compared by Mann-Whitney *U* tests, Kruskal-Wallis tests, or ANOVA (where data were normally distributed, by the Shapiro-Wilk test). Mouse body weight measurements were compared by ANOVA. Gene expression measurements were compared using Kruskal-Wallis tests. Shotgun metagenomics feature count tables were analyzed using R version 4.5.0. and RStudio ^77^. Comparisons of taxon relative abundance were performed at the taxonomic level specified using Wilcoxon rank-sum tests with FDR correction or one-sample Wilcoxon tests using the ggpubr package version 0.6.0 ^93^. Genomes for which taxonomy could not be resolved at the level of analysis (e.g., family) were represented by the lowest rank supported by the reference genome (e.g., order). For CFU, body weight, and gene expression data, a p-value <0.05 was considered statistically significant. For shotgun metagenomics data, an FDR-adjusted p-value <0.1 was considered statistically significant.

## SUPPLEMENTAL INFORMATION

Supplemental Figures 1-6

Supplemental Tables 1,2.

