## Supplemental file for "Targeting catecholate siderophore systems enables precision depletion of gut enterobacteria during colitis"

**SUPPLEMENTAL INFORMATION**

Supplemental Figures 1-6

Supplemental Tables 1,2.

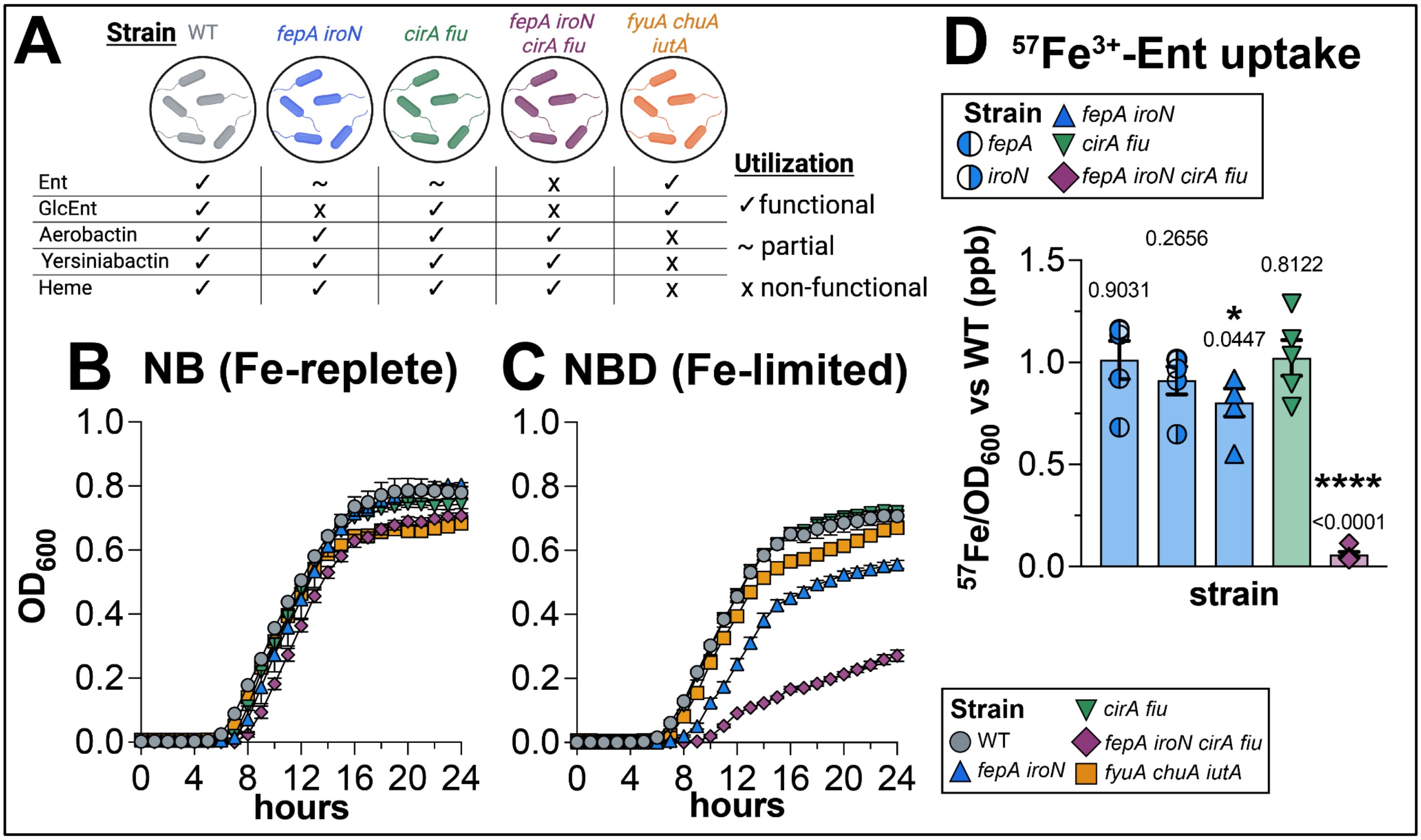


**Suppl. Figure 1.** *In vitro* growth assay comparing AIEC WT and AIEC siderophore uptake mutants in (**A**) Schematic of AIEC siderophore uptake mutants and associated siderophore utilization phenotypes. (**B**) iron-replete conditions (nutrient broth) and (**C**) iron-limited conditions (nutrient broth + 100 μM 2,2'-dipyridyl) by $OD_{600}$ across 24 hours. Points represent mean +/- SEM for n=3 biological replicates per strain. (**D**) Cell-associated $^{57}Fe$ quantified by ICPMS from AIEC after incubation with $^{57}Fe^{3+}$-Ent. Values for siderophore mutants relative to mean AIEC WT and compared by one-sample t-tests.

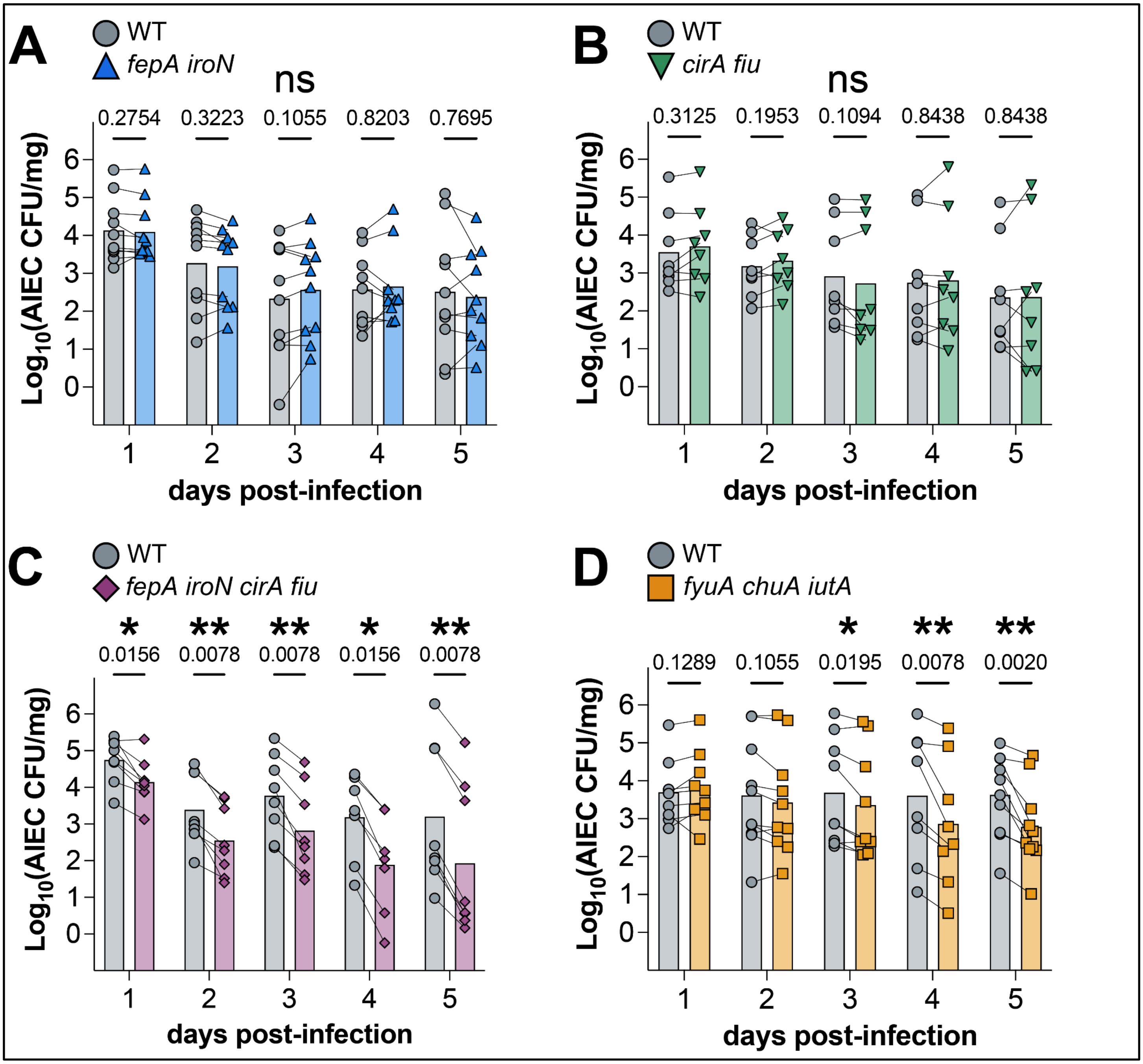


**Suppl. Figure 2.** $Log_{10}$ fecal AIEC CFU/mg in mice co-infected with AIEC WT and (**A**) AIEC *fepA iroN,* (**B**) AIEC *cirA fiu,* (**C**) AIEC *fepA iroN cirA fiu,* or (**D**) AIEC *fyuA chuA iutA*. Bars show mean +/- SEM, points represent CFU measurements for individual animals, and lines connect paired measurements—statistical comparisons made by multiple Wilcoxon signed-rank tests.

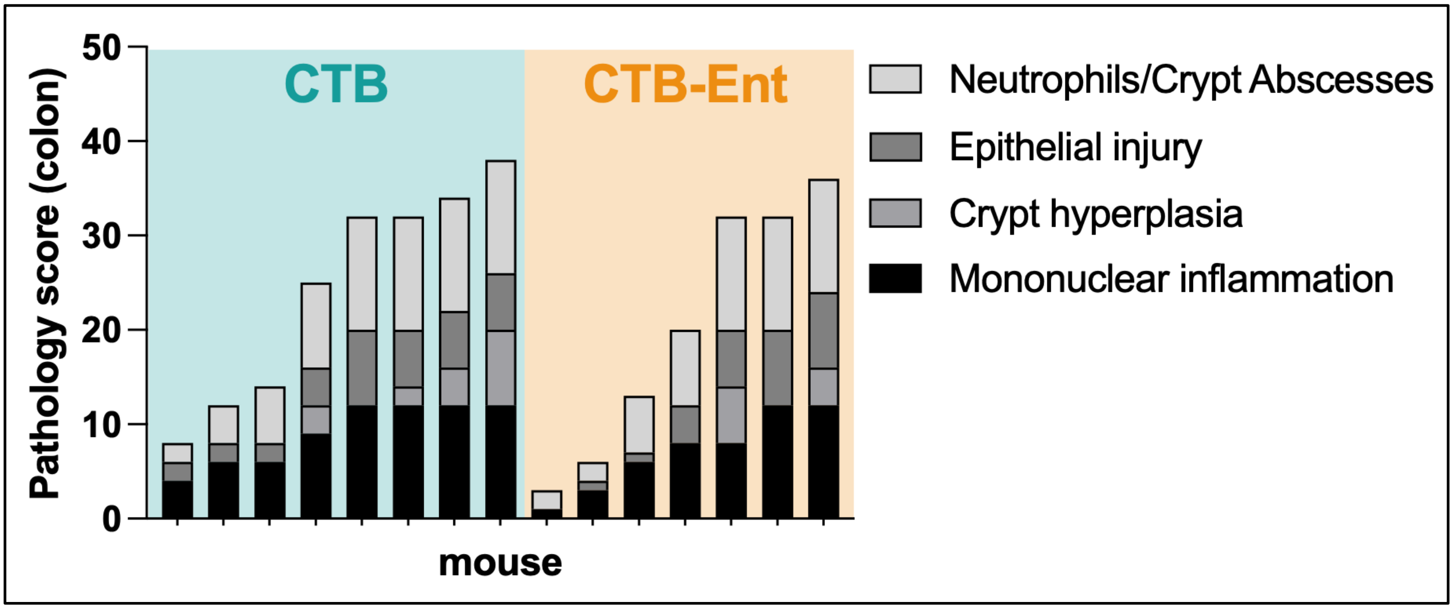


**Suppl. Figure 3.** Distal colon histopathology scores for CTB and CTB-Ent immunized animals.

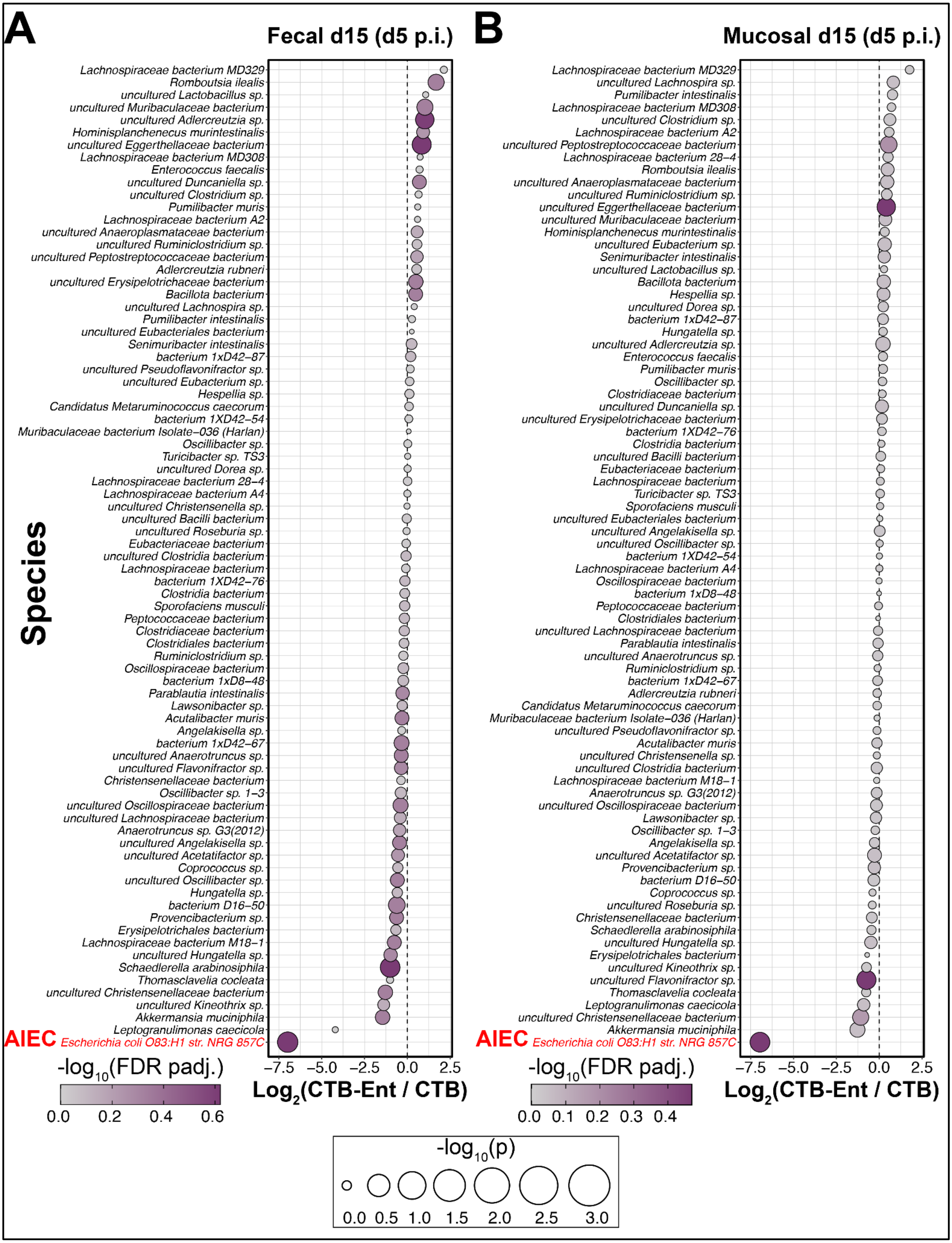

A
Fecal d15 (d5 p.i.)
B
Mucosal d15 (d5 p.i.)
Species
AIEC Escherichia coli O83:H1 str. NRG 857C
-log10(FDR padj.)
Log2(CTB-Ent / CTB)
-log10(p)

**Suppl. Figure 4.** AIEC drives compositional differences between CTB control and CTB-Ent microbiome profiles. (x-axis) $Log_2$-transformed fold change of mean day 15 (day 5 p.i.) CTB-Ent/CTB (y-axis) per-species relative abundances from (**A**) fecal and (**B**) colon mucosal metagenomics profiles. Results of per-species Wilcoxon rank-sum tests are denoted by point size (raw p-value) and color (FDR-adjusted p-value). AIEC fold changes are highlighted in red.

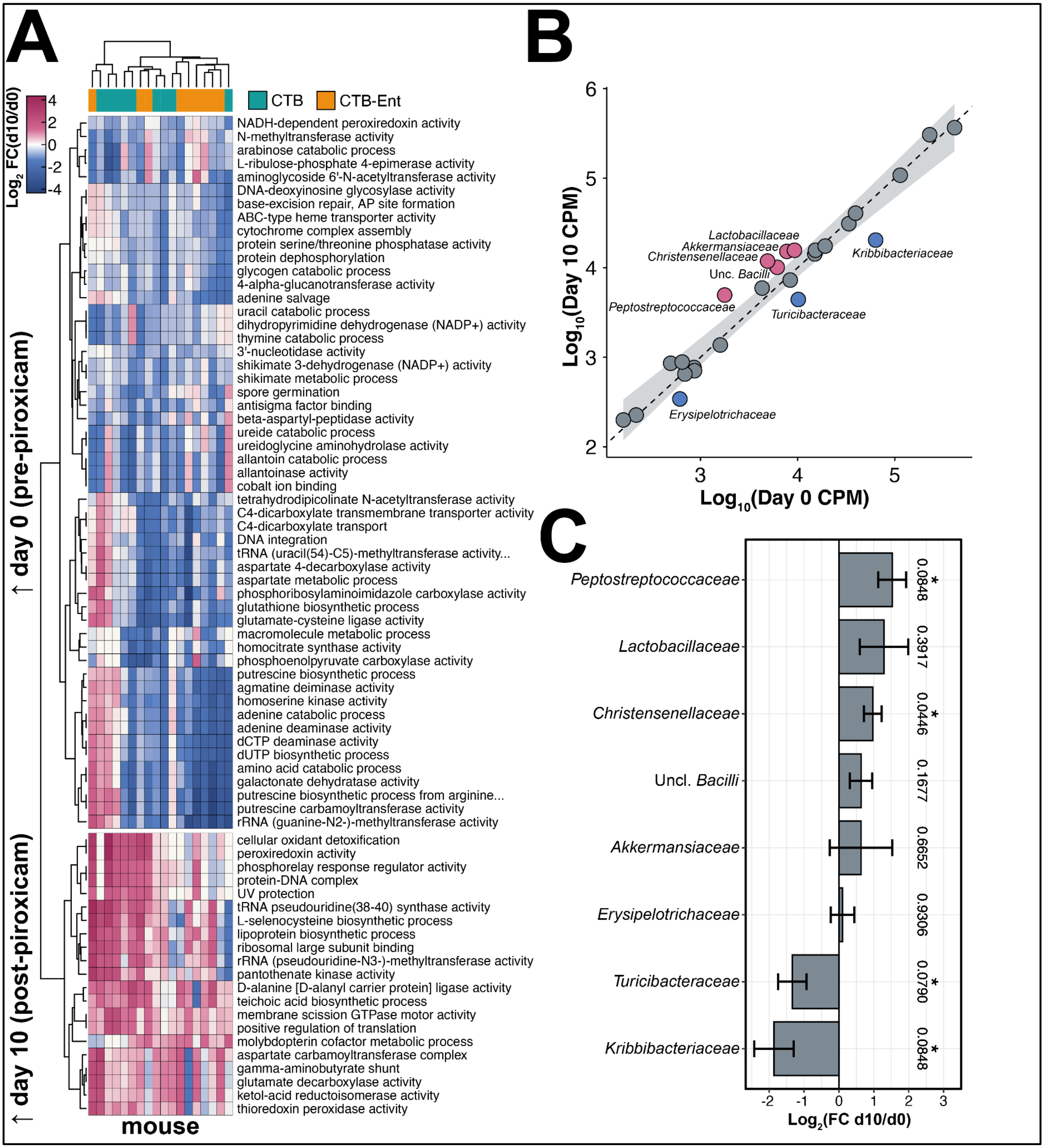


**Suppl. Figure 5. Functional and taxonomic changes in fecal microbiota during piroxicam-accelerated colitis.** Shotgun metagenomics analysis from fecal samples collected before (day 0) and after (day 10) administration of a diet containing 100 ppm piroxicam to *$Il10^{-/-}$* mice. Reads were normalized to counts per million (CPM) and annotated with gene ontology terms (pathways). (**A**) Heatmap of piroxicam-associated pathway changes. Values represent paired $Log_2$ fold change (FC) between samples collected from the same animal day 10 after piroxicam diet administration relative to day 0. Pathways with FDR-adjusted p-values <0.1 and mean |day 10 / day 0|>1.5 are shown. Statistical comparisons made by Wilcoxon signed-rank tests with FDR correction. Columns clustered by pathway abundance and colored annotation bar denote CTB and CTB-Ent immunized mice. (**B**) Linear regressions of (y-axis) day 10 vs (x-axis) day 0 family-level relative abundances. Points represent mean relative abundances and the shaded region represents the 99% confidence interval. Points within the confidence interval are colored in grey. (**C**) $Log_2$ fold change (day 10 / day 0) for bacterial families falling outside the 99% confidence interval in (**B**). Bars represent mean $Log_2$ fold change from paired samples. Statistical significance determined by one-sample Wilcoxon tests with FDR correction. Data represent combined CTB and CTB-Ent immunized mice, as the groups were comparable before AIEC infection.

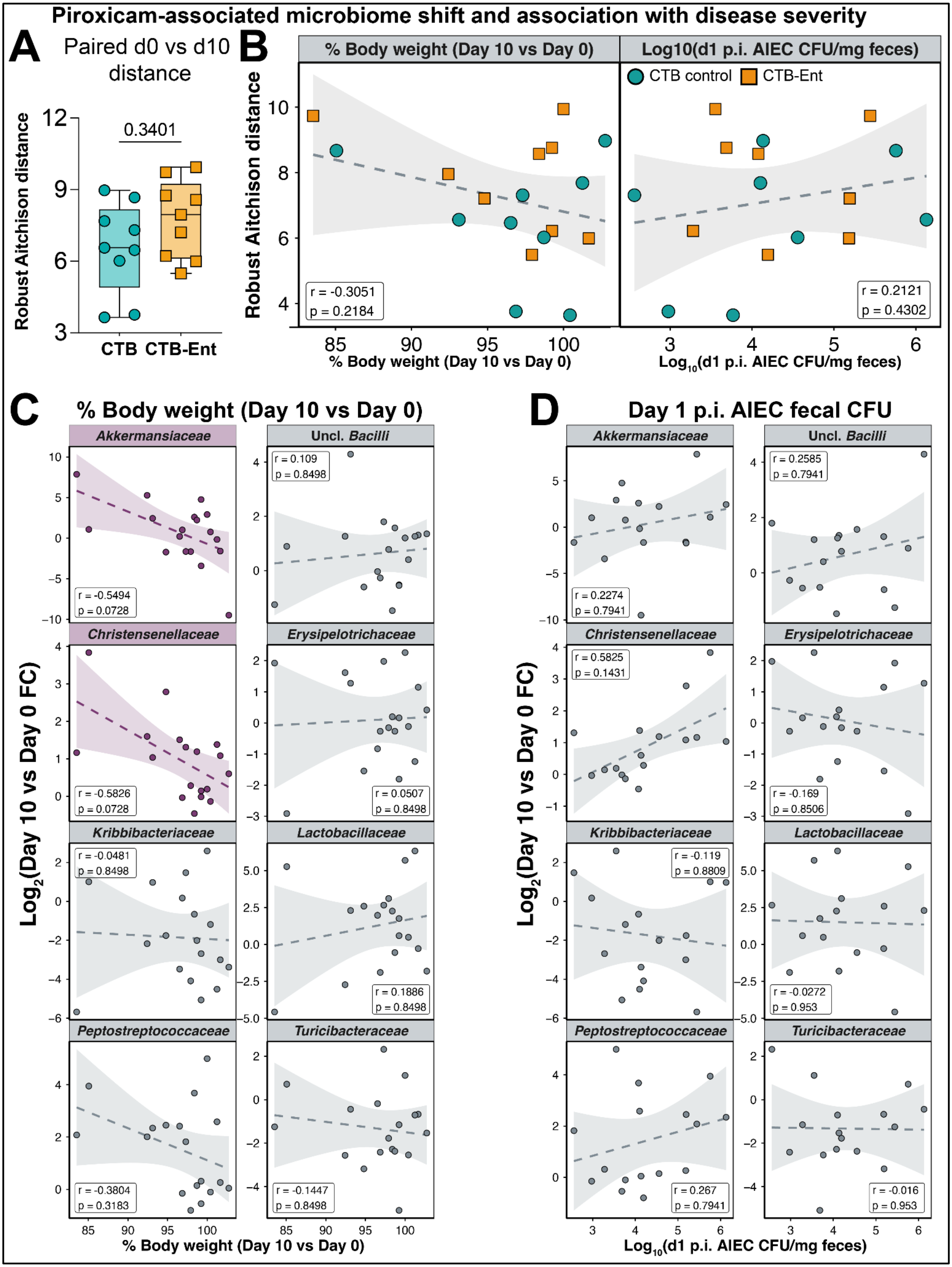

Piroxicam-associated microbiome shift and association with disease severity
A
Paired d0 vs d10 distance
Robust Aitchison distance
0.3401
CTB
CTB-Ent
B
% Body weight (Day 10 vs Day 0)
Log10(d1 p.i. AIEC CFU/mg feces)
CTB control
CTB-Ent
Robust Aitchison distance
r = -0.3051
p = 0.2184
r = 0.2121
p = 0.4302
% Body weight (Day 10 vs Day 0)
Log10(d1 p.i. AIEC CFU/mg feces)
C
% Body weight (Day 10 vs Day 0)
D
Day 1 p.i. AIEC fecal CFU
Akkermansiaceae
Uncl. Bacilli
Christensenellaceae
Erysipelotrichaceae
Kribbibacteriaceae
Lactobacillaceae
Peptostreptococcaceae
Turicibacteraceae
r = -0.5494
p = 0.0728
r = 0.109
p = 0.8498
r = -0.5826
p = 0.0728
r = 0.0507
p = 0.8498
r = -0.0481
p = 0.8498
r = 0.1886
p = 0.8498
r = -0.3804
p = 0.3183
r = -0.1447
p = 0.8498
r = 0.2274
p = 0.7941
r = 0.2585
p = 0.7941
r = 0.5825
p = 0.1431
r = -0.169
p = 0.8506
r = -0.119
p = 0.8809
r = -0.0272
p = 0.953
r = 0.267
p = 0.7941
r = -0.016
p = 0.953
Log2(Day 10 vs Day 0 FC)
% Body weight (Day 10 vs Day 0)
Log10(d1 p.i. AIEC CFU/mg feces)

**Suppl. Figure 6. Association between commensal microbiota shift, body weight change, and AIEC colonization on piroxicam diet (before AIEC infection).** (**A**) Genus-level robust Aitchison distance between paired day 0 and day 10 fecal samples. Points represent individual animals. Statistical comparison made by Wilcoxon rank-sum test. (**B**) (y-axis) Paired robust Aitchison distances between day 0 and day 10 fecal samples plotted against (left) percent body weight change (day 10 after piroxicam diet administration relative to day 0) and (right) day 1 post-infection fecal AIEC CFU/mg. Points represent individual animals. (**C-D**) (y-axis) $Log_2$ fold change (FC) for families with variable relative abundances across piroxicam administration (day 10 vs day 0) in CTB and CTB-Ent t plotted against (x-axis) (**C**) percent body weight (day 10 after piroxicam diet administration relative to day 0) or (**D**) day 1 post-infection fecal AIEC CFU/mg. Points represent fold change values in paired day 10 and day 0 samples from the same animal. Lines represent linear regression and shaded regions denote 95% confidence intervals. Statistics from correlation analysis; r = Pearson's R and p = FDR-adjusted p-value. Data represent combined CTB and CTB-Ent immunized animals, as the groups were comparable before AIEC infection.

**Supplementary Table 1**

| **Strain or plasmid** | **Relevant characteristics** | **Source or Reference** |
|---|---|---|
| GN0200 | *Escherichia coli* AIEC O83:H1 NRG857c wild-type | 33 |
| GN0207 | *Escherichia coli* AIEC O83:H1 NRG857c Δ*iroN*::GmR | This study |
| GN0215 | *Escherichia coli* AIEC O83:H1 NRG857c Δ*fepA*::GmR | This study |
| GN0224 | *Escherichia coli* AIEC O83:H1 NRG857c Δ*iroN*::FRT Δ*fepA*::GmR | This study |
| GN0247 | *Escherichia coli* AIEC O83:H1 NRG857c Δ*cirA*::FRT Δ*fiu*::GmR | This study |
| GN0241 | *Escherichia coli* AIEC O83:H1 NRG857c Δ*iroN*::FRT Δ*fepA*::FRT Δ*cirA*::FRT Δ*fiu*::GmR | This study |
| GN0255 | *Escherichia coli* AIEC O83:H1 NRG857c Δ*fyuA*::FRT Δ*chuA*::FRT Δ*iutA*::GmR | This study |
| GN2600 | *Akkermansia muciniphila* BAA-835 | 74 |
| GN2800 | *Limosilactobacillus reuteri* mouse isolate | This study |
| pGmCDF_Frt | Template plasmid for amplifying the KO product with a gentamicin-resistance cassette. Plasmid also has spectinomycin resistance cassette. | 76 |
| pKD46_Km | Plasmid expressing Red recombinase. Arabinose-inducible and temperature-sensitive (propagate at 30˚C). | 76 |
| pCP20_Km | Plasmid expressing the FLP recombinase to flip out the gentamicin-resistance cassette. Temperature sensitive. | 76 |

**Supplementary Table 2**

| Name | Sequence (5’ - 3’) | Source |
|---|---|---|
| iroN_KO_1_F | GGTATGAGAATTAACAAAATCCTCTGGTCGCTAACTGTGCTCCTGTAACAATTCCCCTGTAGAAATAAT | This study |
| iroN_KO_1_R | CTACGATCAGAATGATGCGGTAACTCCGGCATAGTAAGCCCGGGTGATGTGCCATGGGAAGT | This study |
| iroN_external_1_F | TCTGGCAAGGATGTGAGCTT | This study |
| iroN_external_1_R | GTTCAATGCCTCCTTTCGCC | This study |
| iroN_int_1_F | CAGATTCTCGATTTCGAAGCC | This study |
| iroN_int_1_R | CTGCGAAAGATTCAGACTGG | This study |
| FepA_KO_2_F | GCGACCATGCCCGACAGTTGCAATTCGTGGCAAAAATGCAGGAAAAAACATAACAATTCCCCTGTAGAAATAA | This study |
| FepA_KO_2_R | CATCCGGCATGAACTACGCGCACTTTGTCAACAATCTGACGTTAGCTTCAATCATTCGAACCGGTACCCA | This study |
| fepA_external_1_F | AATTAATCAGCTGCACTGCG | This study |
| fepA_external_1_R | AGGTATGCGTAGTTTTCATGTCA | This study |
| FepA_int_1_F | AGAACACCAACTCTGACG | This study |
| FepA_int_1_R | GCCGACAATACTGTGATG | This study |
| cirA_KO_1_F | CTGCCGTACGCAAGGGGACGTAAAGAAGATGTGAGCGATACCCATTAACAATTCCCCTGTAGAAATAAT | This study |
| cirA_KO_1_R | TAGAGGATCACCTGCGCTTCCATGCGCAGTATTTACTGAAGTGAAAGTCCGGGTGATGTGCCATGGGAAGT | This study |
| cirA_external_1_F | CTGTTCCGGCTTTCTGGGAT | This study |
| cirA_external_1_R | CTTGCTGACCAACTGCAACC | This study |

| | | |
|---|---|---|
| cirA_int_1_F | CGTCACGACAAACTTAGC | This study |
| cirA_int_1_R | GATCGACAGTTTCCATTCATCG | This study |
| fiu_KO_F | CGGGATTAACAGTGGCATCGC ATCCGCAGAGATGCTTTCTCGT GACAGTGTAACAATTCCCCTGT AGAAATAA | This study |
| fiu_KO_R | GGTGATAACGGTAGCCGCTCTT ATTGATTGAGGCGACGTAATCG GTATCAATCATTCGAACCGGTA CCCA | This study |
| fiu_ext_F | CCTAAGCGATCTGGAGGATAAA C | This study |
| fiu_ext_R | CTGCTCTCGCCATCAACATTAG | This study |
| fiu_int_F | TGGATAAACGCCTGCTACTTAC | This study |
| fiu_int_R | AGATTGCGATTAACCCGATACC | This study |
| fyuA_KO_1_F | GTCGGTATTTTTTGGCGTTTCG CCGTCTTACAGGGACTCACAAC ATAACAATTCCCCTGTAGAAAT AAT | This study |
| fyuA_KO_1_R | GATACCTTTTCGGTATCCCTTTT ACAATAATCAGAAGAAATCGGT GATGTGCCATGGGAAGT | This study |
| fyuA_external_1_F | GAGCGGATTTGCGCGTTTAT | This study |
| fyuA_external_1_R | TGTACAGGCCTATGGGGTGA | This study |
| fyuA_int_F | TTTCGGGTAGCACCACCATC | This study |
| fyuA_int_R | CGATAACTGTCGCGGCTACT | This study |
| chuA_KO_1_F | GTATTAACCCAGGCAGCCAGCA TTTTTTATTTTCCTCATGGAGAA TCGCTTAACAATTCCCCTGTAG AAATAA | This study |
| chuA_KO_1_R | GCGGCGACCAGTACTCTTTATC GAAGGCGTTGCCCAATACCAG AGTAGTGATCATTCGAACCGGT ACCCA | This study |
| chuA_ext_F | TACGCTAATAGCTTCCCG | This study |
| chuA_ext_R | CTCTCTTCCTTCCAGATAAATG C | This study |
| chuA_int_F | CGATTGGTCGCTTCTATACC | This study |

| | | |
|---|---|---|
| chuA_int_R | CTGCTGCCCTTGATAACTG | This study |
| iutA_KO_1_F | GGAGGCTGCTTCTTTAACTCGCTACACAGCATCTTTGGGCTGATTTTAACAATTCCCCTGTAGAAATAA | This study |
| iutA_KO_1_R | GGAAGCTGATAACTGCCGAGCAGATCCACGGTGGTATAGCCATCGACCATCATTCGAACCGGTACCCA | This study |
| iutA_ext_F | GCGTAATGGGACGTGATTTATTC | This study |
| iutA_ext_R | GGTCAGAACAGCACAGAGTAG | This study |
| iutA_int_F | GTCTTTCCGTGGCATATCAG | This study |
| iutA_int_R | GTTTGCTGTTCAGAGTCAGTTTC | This study |
| GmR_internal_1_R | CCCGCATGGATTTGACTTGG | This study |
| GmR_internal_2_F | GCAGGTCGACTCTAGAGGAT | This study |
| GmR_int_3_R | CAAGGTTCTGGACCAGTTGC | This study |
| GmR_int_4_F | ACTCCGATTACCTCGGGAAC | This study |
